## Supplementary figures and images for "Spatiotemporal brain complexity quantifies consciousness outside of perturbation paradigms"

### Supplementary Figure S1

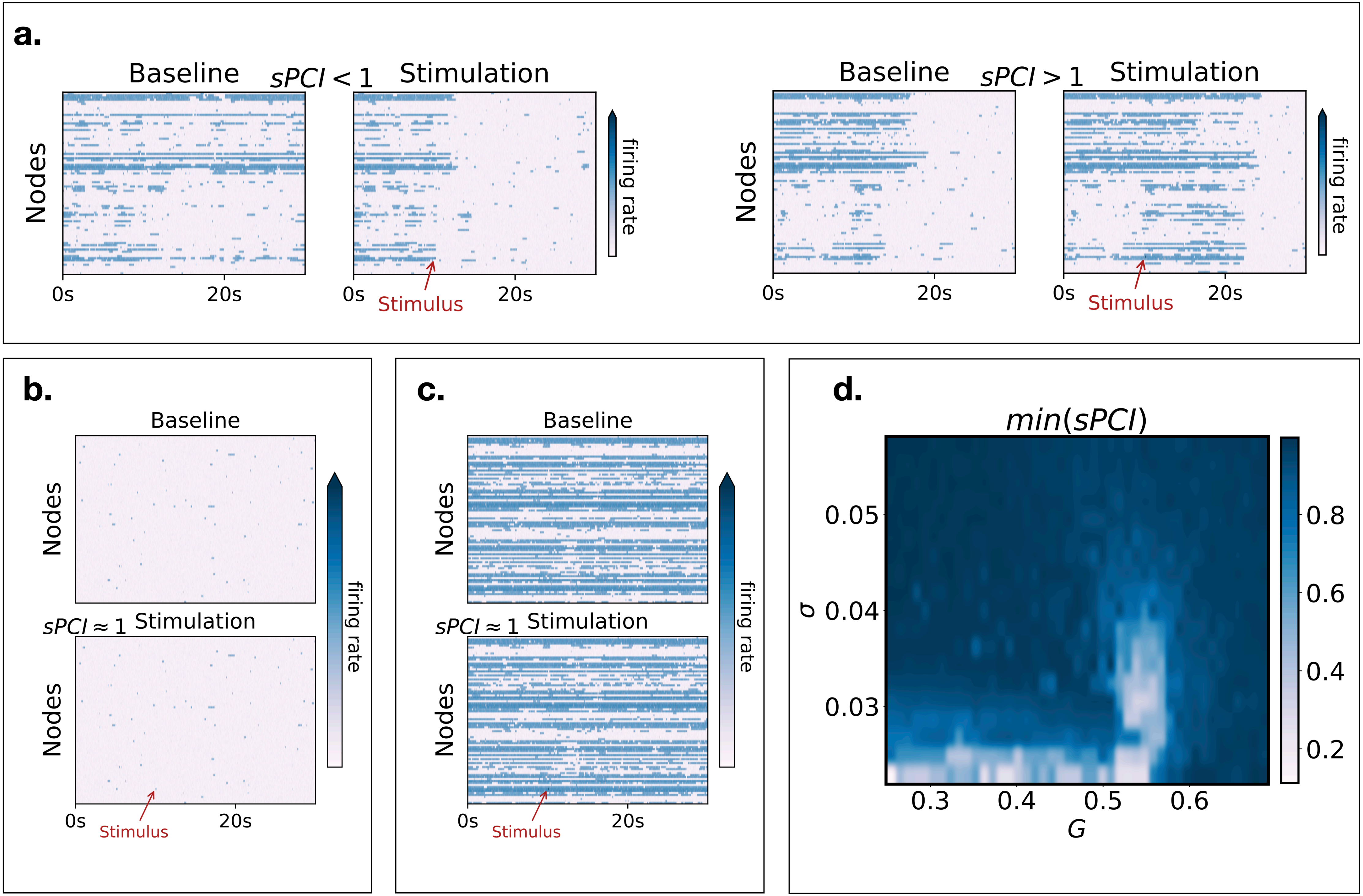
